# Mathematical model of molecular evolution through a stochastic analysis

**DOI:** 10.1101/699264

**Authors:** Joel Valdivia Ortega

**Affiliations:** Universidad Nacional Autonoma de Mexico

## Abstract

Today we are able to describe genome evolution but, there are still several open questions on this area such as: Which is the probability that a mutation occurs at a nucleotide level? Is it possible to predict the evolution of a particular genome?, or talking about preservation, is there a way to simulate the genetic diversity for endangered species? In this paper it is shown that it is possible to make a mathematical model not only of mutations on the genome of species, but of evolution itself, including factors such as artificial and natural selection. It is also presented the algorithm to obtain the probabilities of mutation for each specific part of the genome and for each specie.

Even more, it is presented a mathematical method to estimate the amount of generations between two related genomes and a function capable of predict the amount of mutations through time a genome will suffer.

The potential of having this tool is giantic going from genetic engineering applied to medicine to filling up blank spaces in phylogenetic studies or preservation of endangered species due to genetic diversity.

## 2 Introduction

This paper is to present a mathematical model of molecular evolution, letting us recreate the process in which one genome became another. In order to do so, we will use a stochastic method and after that we well explore some of its implications.

For definition, a discrete Markov chain [1] is a stochastic model consisting on a discrete random variable *x_n_* that evolves with some parameter *n* such that the probability of each state depends only on the last state, i.e.

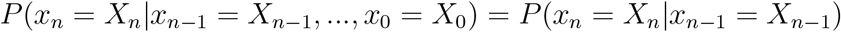

This mathematical tool has the property that the evolution of the whole system can be described by the *transition matrix P*, which contains the probabilities of going from one state to another, meaning that the entrance of the *i*-th row and *j*-th column is *P_ij_* = *P*(*x_n_* = *X_i_*|*x*_*n*−1_ = *X_j_*). This us very helpfull because if we are interested in the probability distribution the system will follow after *k* steps, all we have to do is calculate *P^k^*.

Since the mutations on molecular evolution happen randomly, we are able to describe each nucleotide on the genome as a random variable which varies through generations and has the DNA nucleotides as its range. We will represent them as *x_n_* ∈ {*A, C, G, T*}, where *A* stands for adenine, *C* for cytosine, *G* for guanine and *T* for thymine. Even more, since we know that the mutations on parents do not influence the mutations on the children, we have that the molecular evolution behaves as a discrete Markov chain, so now we are interested in getting the transition matrix to describe the system.

In order make it easier, we will assume the probabilities of mutation do not vary between small pieces of the genome so we would be able to describe a portion of 10000 nucleotides with the same transition matrix.

In the next sections, we will analyze a chain of 10000 nucleotides corresponding to genomes of a wolf and several dogs. We chose this genomes as an example because the genetic relation between this species has been proved through several methodes.

## 3 Stochastic analysis of *canis lupus* and *canis lupus familiaris* genomes

The chosen genomic chains are the thousand nucleotides taken from the National Center for Biotechnology Information (NCBI) and correspond to a wolf [2], a boxer [3], a poodle [4], a yorkshire [5] and a beagle [6].

Let *w*(*m*) be the *m*-th nucleotide of the wolf’s genome and *d*(*m*) the *m*-th nucleotide of a dog’s genome such that *d* ∈ {*boxer, poodle, yorkshire, beagle*} =: *D*.

What we will do now is obtain the transition matrix *P*′, so we will define our space of states as the ordered set *N* = {*N*_1_ = *A*, *N*_2_ = *C*, *N*_3_ = *G*, *N*_4_ = *T*} and we will define

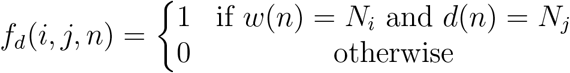

where *d* ∈ *D*, so we can write *P*′ in such a way that 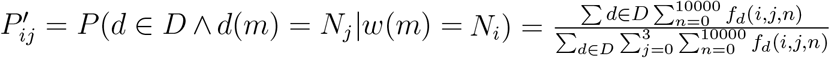 which corresponds to the probability that if a nucleotide on the wolf’s genome if *N_i_*, on a dog’s genome it becomes a *N_j_*.

According to The Smithsonian [7], the domestication of the wolf occured about 30000 years ago and since then the evolution of the dog began. Supposing a new generation of dogs is born each year on average, we affirm that if *P* is the transition matrix for the nucleotides, *P*′ = *P*^30000^, realizing it is a matrix 4 × 4.

In order to obtain *P*, let us define

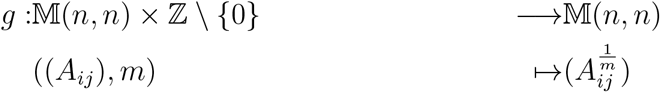

and if *x* ∈ ℂ, we denote 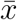 as its complex conjugate and define

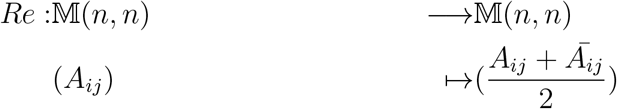

so if *P*″ is the matrix which has the eigen values of *P*′ on its diagonal and 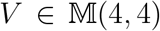 such that *P*′ = *VP*″*V*^−1^, therefore we may obtain *P* = *Re*(*V_g_*(*P*″, 30000)*V*^−1^) because of the behaviour of diagonal matrices under multiplication and group theory. In our example it corresponds to the next matrix.

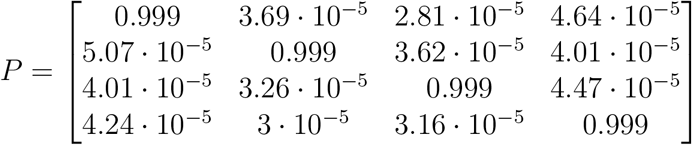

The previous matrix can be represented as a directed graph in which the vertex are the nucleotides and the values of the edges correspond to the probability of transition.

Let us realize that this diagram matches the daily experience, since it does not allow several mutation from one generation to another. On the other hand it also corresponds to the observed characteristics of evolution, for example the fact that no edge has a null value means that molecular evolution cannot be separated into several Markov chains while it also let us affirm that molecular evolution will never arrive to an equilibrium point nor finish.

## 4 Testing the transition matrix

Now we have our transition matrix, if we claim molecular evolution can be described with a Markov chain, the less we should be capable of doing is begin with the wolf’s genome and arrive to each dog’s genome.

In order to do this we wrote an *artificial selection* program which, beginning with the genome of the wolf and choosing a specific dog as a target, simulated 50 children and chose the 2 of them which were more similar to the dog. After doing that, the program simulated the chosen children to have a breed between themselves and chose the specimen which had the closest genome to the dog we were attempting to reach, simulating now that it had 50 children applying this process iteratively.

The process was made for all the four dogs and in every case the program converged but since this is a probabilistic method, now we must place the question: does it has something do to with the dogs or we are just reaching the desired genome for the way the algorithm was made?

**Figure 1:**
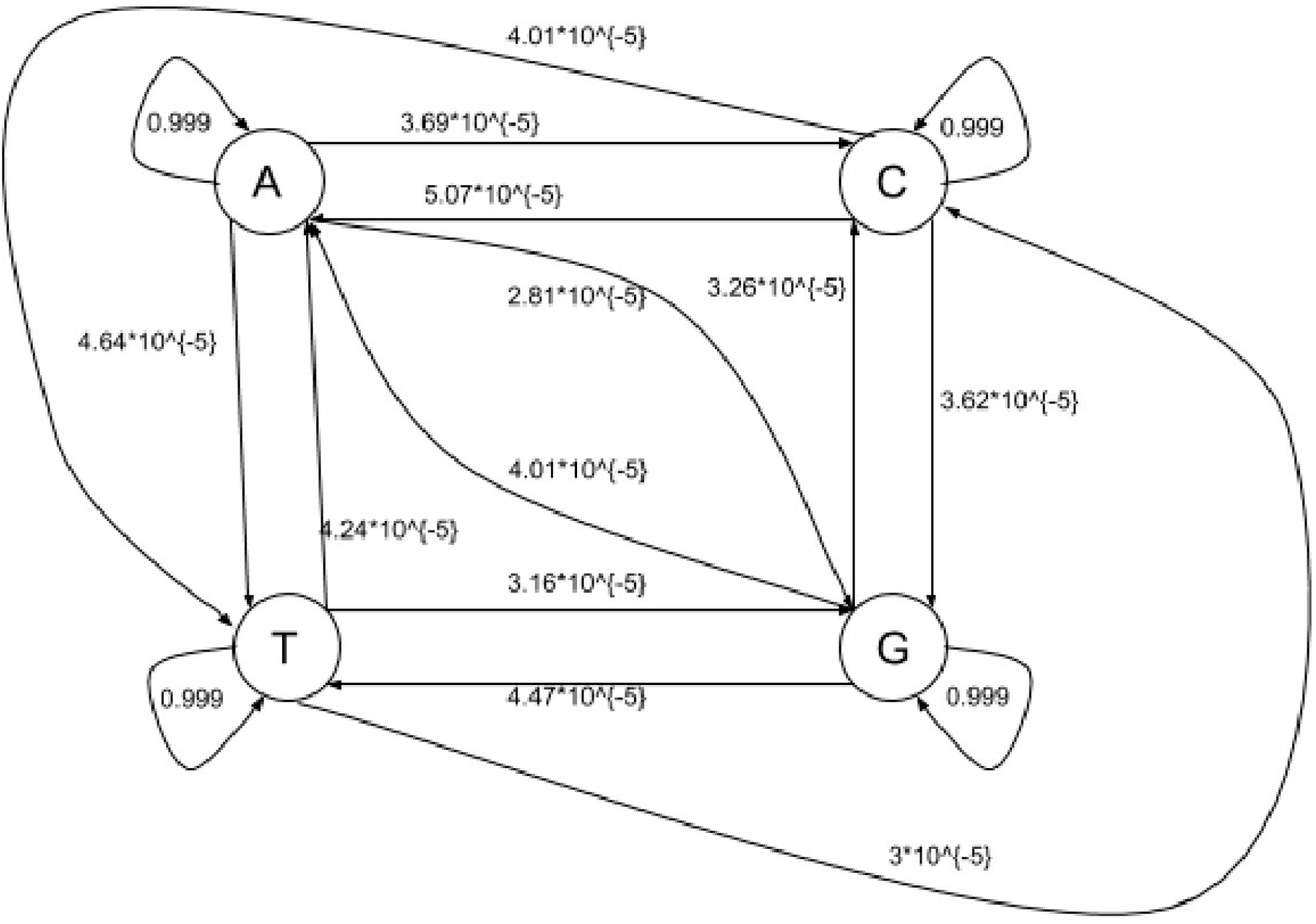
Diagram of the system’s transition probabilities

To answer this, we chose a randomly generated genome as the target and ran the program in order to see if the program also converges in this case and if so, if there was a qualitative or quantitative difference between the other cases. The results of simulating 100 times the process are shown below.

As we can see on Table 1, there is no significant difference between the results from the dogs and the results from the randomly generated genome so this means that this artificial selection program is not good to predict the genome’s evolution outside the known species.

**Table 1:** Statistical data about the results of computational process to reach the dog’s genome and the randomly generated genome

|  | Boxer | Poodle | Beagle | Yorkshire | Random genome |
| --- | --- | --- | --- | --- | --- |
| Average amount of generations taken to reach the genome | 1036.434 | 996.94 | 1042.538 | 1062.708 | 1077.428 |
| Standard deviation of the results between themselves | 135.023 | 127.864 | 137.112 | 135.142 | 134.419 |
| Standard deviation as a percentage of the average | 13.027 | 12.825 | 13.151 | 12.716 | 12.475 |

However, the fact that the program drive us with probabilistic methods from the wolf’s genome to each of the dog’s genome and due to the law of large numbers, we can affirm that this artificial selection program give us the most probable molecular evolution between the known species, letting us know the “genomic history” that connects one specie with the other.

But we are trying to design a way to study the whole process of evolution, not just an interpolation between species so what we made was, using our transition matrix and for computational simplicity selecting just the first n nucleotides from each dog’s genome, to simulate 7 probable parents for each of our 4 dogs and then to choose the closest ones between each other. After that, we iterated the simulation for the selected parents and ran the program until it converges with a difference between the genome from the parents of 1% of the total amount of nucleotides.

Once we arrived to this difference, we calculated the percentage of differences between the genome we arrived to and the wolf’s genome. Given a n, we repeated this process 20 times for statistical purposes and we called *discordance of n D_n_* to the average this quantities.

On the other hand and keeping the given n, we calculated the average percentage of similarity between the chain of nucleotides of each dog and the one from the wolf and to that number we called *concordance of n C_n_*.

It is worthy to remark that, since we did not use directly the wolf’s genome in this algorithm but to get the transition matrix, whatever the result is, it will not depend on anything but the begining genomes and the transition matrix.

We took 74 different n’s represented on Figure 2 and the result was that the average sum of the concordance and the discordance of each n *C_n_* + *D_n_* is 100.03% with a standard deviation of 2.81%.

**Figure 2:**
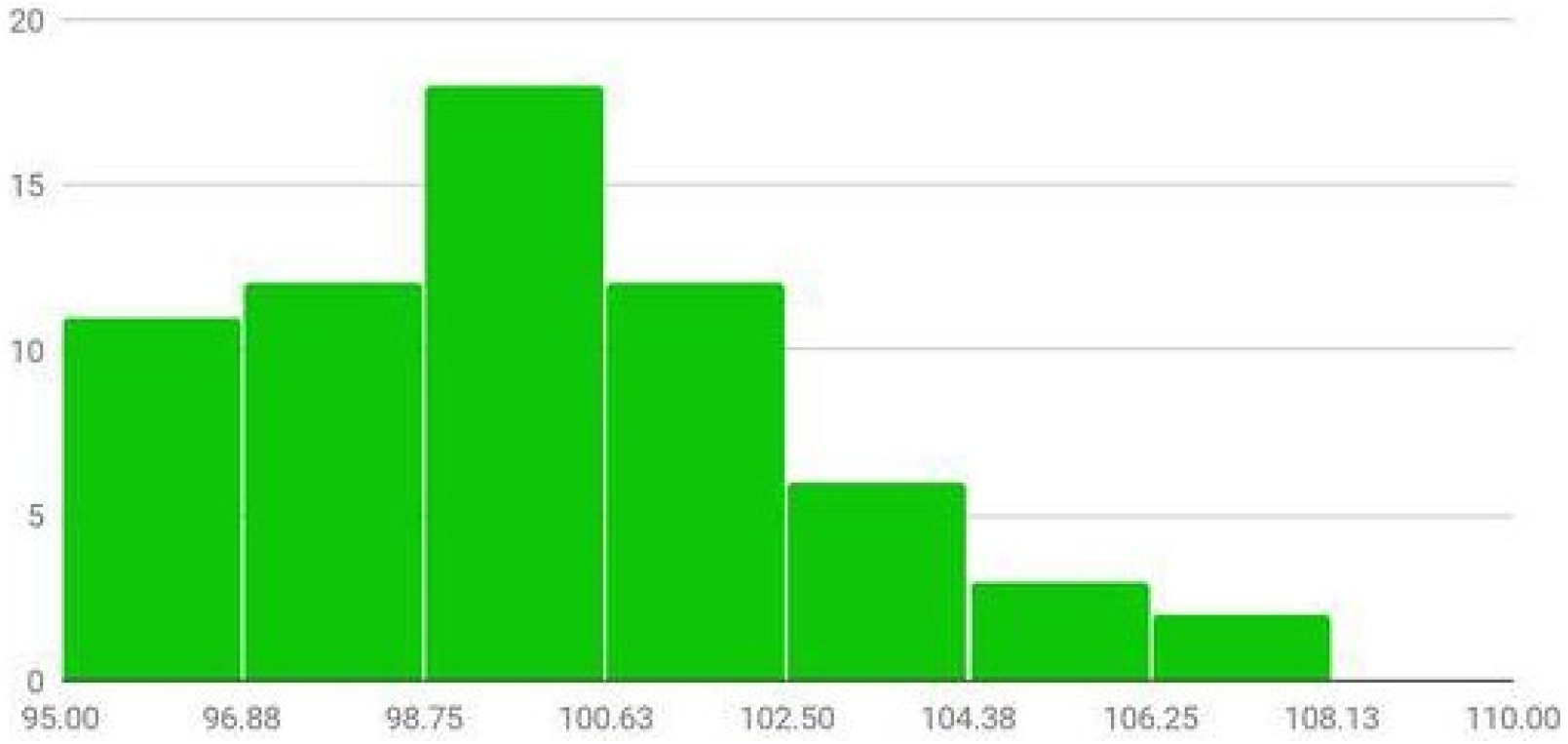
Histogram of *C_n_* + *D_n_*

We may take this as a truth for any number of nucleotides because of the next reason: given an enumeration of the genome’s nucleotides, let us take the m-th one such that *w*(*m*) = *N*′ and *d_j_*(*m*) = *N_j_* where 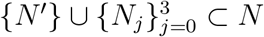 and *d_j_* ∈ *D*, ∀*_j_* ∈ {0,…, 3}.

Following our algorithm we will apply the transition matrix to *d_j_* until *d_j_* = *d_i_*, ∀{*i, j*} ⊂ {0,…, 3} and in our example we have that the probability for *d_j_* to change has the order of 10^-4^, so on average we will need 10^4^ iterations for the nucleotide to change. In other words, the distribution of probability that *d_j_* will obey is given by *P*^10^4^^ ≈ *P*^30000^, i.e. the distribution corresponding to the wolf’s genome.

This result is the final confirmation that modelling molecular evolution as a discrete Markov chain is accurate, that our transition matrix has the correct probabilities of transition between nucleotides and that we can not only interpolate but also extrapolate predictions of the genome’s evolution through given species because we know that the bigger the chain of DNA we take, the more similar the dog’s genome will be to the wolf’s one and therefore the discordance will tend to zero, i.e. *C_n_* + *D_n_* = 100 ⟹ *C_n_* = 100 − *D_n_* ⟹ lim_*n*→∞_ *C_n_* = 100 − lim_*n*→∞_ *D_n_*, but from statistical biology [8] we know that lim_*n*→∞_ *C_n_* = 100 ⟹ lim_*n*→∞_ *D_n_* = 0.

In other words, the probability that the resulting genome of this algorithm converges to the wolf’s genome tends to 1 as we take more nucleotides in our genome, or written with mathematical symbols *lim*_*n*→∞_*P*(resulting genome = wolf’s genome) = 1.

## 5 Measuring the distance between two related genomes

Now that we have our transition matrix and that we have confirmed its validity, we are able to describe the behavior of the system and make computational experiments.

For that, let be *q_n_* an individual which genome is the dictated by the probability distribution *P^n^w*(*m*) and let be 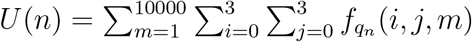, i.e. the number of identical nucleotides between *q_n_* and *w*.

We will now search for a differential equation for *U* and to do so we will take the folliwing hypothesis:

1. The number of new differences depends on the quantity of equal nucelotides on the genome, then 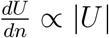.
2. The probability of a mutation on a specific entry of the genome does not depend on the number of generations without a change, which means 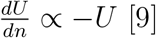 [9].
3. It is possible that a nucleotide from *q_n_* mutates in such a way that it becomes the same as in *w*, therefore 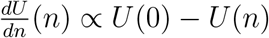.

So we propose the next differential equation

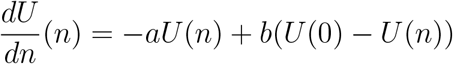

which general solution, taking *n* as a continuous variable, is 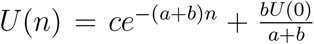 and taking the condition *q*_0_ = *w*, we have

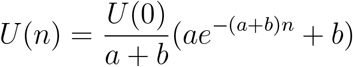

where *a* represents the probability that if *q_n_*(*m*) = *w*(*m*), *q*_*n*+1_(*m*) ≠ *w*(*m*) for any nucleotide *m* and *b* represents the probability that if *q_n_*(*m*) ≠ *w*(*m*), *q*_*n*+1_ = *w*(*m*).

We will obtain those probabilities from the transition matrix as 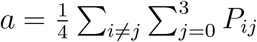 and 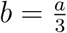 inspired from the interpretation of the entrances of *P*, so we get

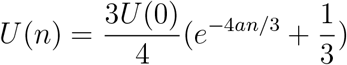

where we can see that *q_n_* will approach asymptotically to a randomly generated genome, explaining the genetic diversity in nature.

In our example *U*(0) = 10000 so putting the numeric values we get

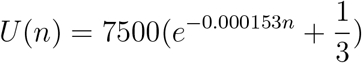

By overlapping the graphs from *U*(*n*) and a computational simulation we obtain the following figure.

Because of the factor of correlation we can affirm that this equation represents correctly the phenomena which means that we are able, for example, to calculate the number of generations between two given genomes if they have a phylogenetic relation by the following the general equation:

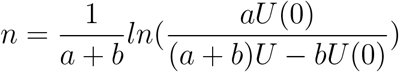

This equation gives an average distance to our example of 34054 ± 70 generations between the dogs and the wolf, which is a very good result we built the transition matrix under the supposition that the number of generations between the dog and the wolf is 30000.

Let us notice that this method can be generalized to any specie since we did not use a specific characteristic from the dogs neither from the wolf but to have a phylogenetic relationship.

**Figure 3:**
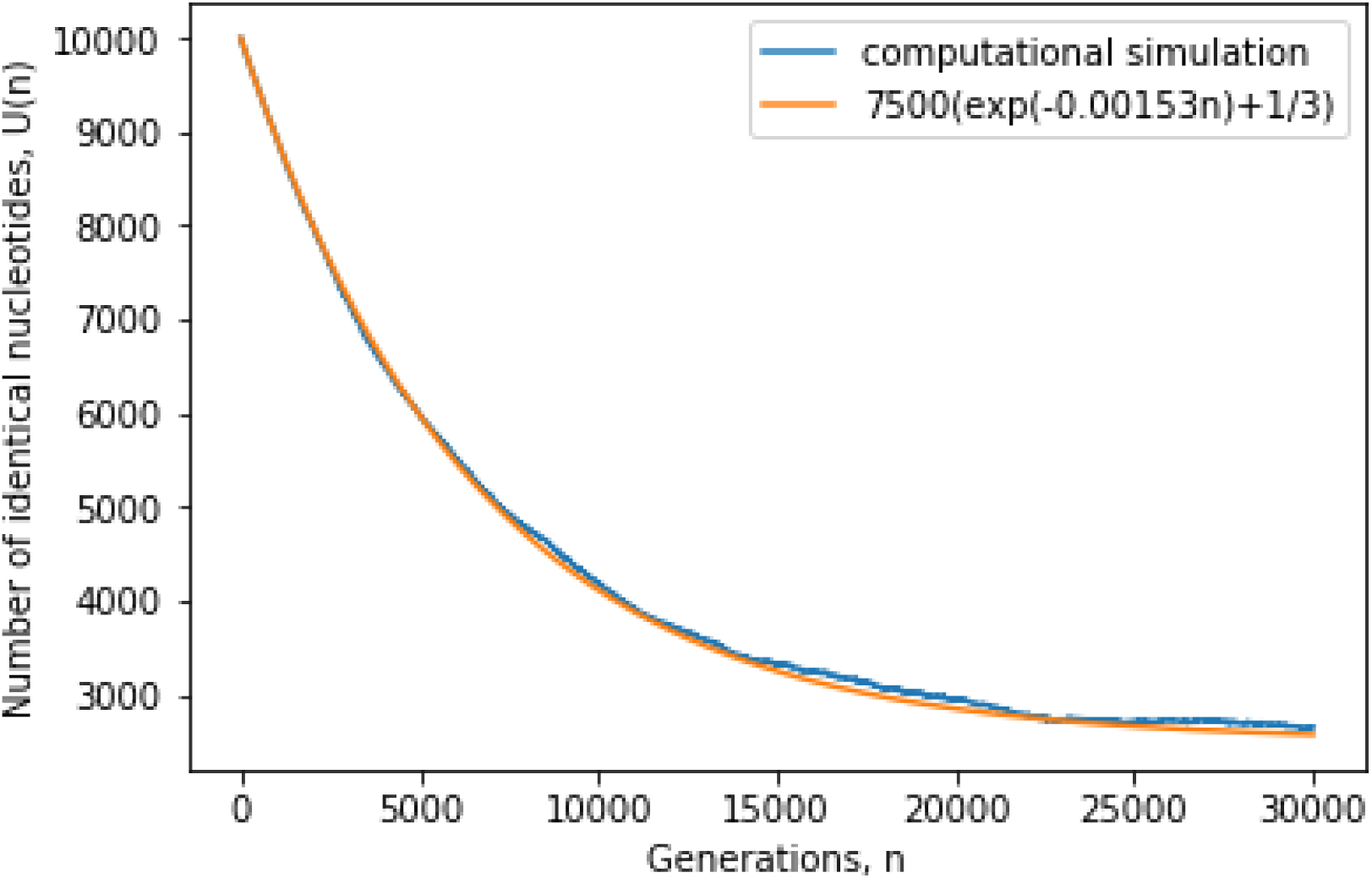
Graphs from a computational simulation and *U* with a factor of correlation *R*^2^ = 0.9996

